# A balance of outward and linear inward ionic currents is required for the generation of slow wave oscillations

**DOI:** 10.1101/136887

**Authors:** Jorge Golowasch, Amitabha Bose, Yinzheng Guan, Dalia Salloum, Andrea Roeser, Farzan Nadim

**Author notes:** **Corresponding Author:** Jorge Golowasch Federated Department of Biological Sciences, NJIT, 100 Summit St, CKB 337, University Heights, NJ 07103.

## Abstract

Regenerative inward currents help produce slow oscillations through a negative-slope conductance region of their current-voltage relationship that is well approximated by a linear negative conductance. We used dynamic clamp injections of a linear current with this conductance, *I_NL_*, to explore why some neurons can generate intrinsic slow oscillations whereas others cannot. We addressed this question, in synaptically isolated neurons of the crab *Cancer borealis*, after blocking action potentials. The pyloric network consists of distinct pacemaker group and follower neurons, all of which express the same complement of ionic currents. When the pyloric dilator (PD) neuron, a member of the pacemaker group, was injected with *I_NL_* using dynamic clamp, it consistently produced slow oscillations. In contrast, the lateral pyloric (LP) or ventral pyloric (VD) follower neurons, failed to oscillate with *I_NL_*. To understand these distinct behaviors, we compared outward current levels of PD, LP and VD neurons. We found that LP and VD neurons had significantly larger high-threshold potassium currents (*I_HTK_*) than PD, and LP had lower transient potassium current, *I_A_*. Reducing *I_HTK_* pharmacologically enabled both LP and VD neurons to produce oscillations with *I_NL_*, whereas modifying *I_A_* levels did not affect *I_NL_*-induced oscillations. Using phase-plane and bifurcation analysis of a simplified model cell, we demonstrate that large levels of *I_HTK_* can block *I_NL_*-induced oscillatory activity, whereas generation of oscillations is almost independent of *I_A_* levels. These results demonstrate the importance of a balance between inward pacemaking currents and high-threshold K^+^current levels in determining slow oscillatory activity.

## Introduction

Leak currents are key determinants of neuronal excitability (Brickley et al. 2007; Lu and Feng 2012; Lutas et al. 2016; Rekling et al. 2000) and can be regulated by many different neuromodulators, which can modify the activity to either silence neurons, or to induce spiking or oscillatory activity (Bayliss et al. 1992; Cymbalyuk et al. 2002; Egorov et al. 2002; Lu and Feng 2012; Lutas et al. 2016; Talley et al. 2000; van den Top et al. 2004; Vandermaelen and Aghajanian 1983; Xu et al. 2009). Leak currents have been proposed to control pacemaker rhythm generation (Amarillo et al. 2014; Blethyn et al. 2006; Cymbalyuk et al. 2002; Koizumi and Smith 2008; Lu and Feng 2012; Pang et al. 2009; Yamada-Hanff and Bean 2013; Zhao et al. 2010). Often, leak currents act to bring the membrane potential to within a range of voltage where other currents can activate and produce a new state of activity (Brickley et al. 2007; Lu and Feng 2012; Rekling et al. 2000; Yamada-Hanff and Bean 2013).

Regenerative inward currents, such as persistent Na^+^- and low-threshold Ca^++^-currents, are essential for the generation of oscillatory neuronal activity (Amarillo et al. 2014; Del Negro et al. 2002; Dunmyre et al. 2011; Jahnsen and Llinas 1984; McCormick and Huguenard 1992; Yamada-Hanff and Bean 2013; Zhao et al. 2010). These currents can be divided into two almost linear components, only one of which is sufficient and necessary to generate the oscillations (Bose et al. 2014; Zhao et al. 2010). Since this component is linear, we refer to it as a leak current; however, it has negative slope conductance (hence, a negative-conductance leak current, *I_NL_*). It is a leak current in the sense that, when combined with the standard leak current (*I_L_*), the total current *I_NL_*+*I_L_* remains linear, and yet it is a key determinant of neuronal excitability. The mechanism by which *I_NL_* controls oscillatory activity is by destabilizing the resting state of the cell (Bose et al. 2014) thereby increasing the voltage of the cell to a point where outward currents can turn on and bring the voltage back to hyperpolarized levels. In this way, when *I_NL_* dominates over *I_L_*, the total linear current can be a *de facto* pacemaker current (Bose et al. 2014; Zhao et al. 2010).

Outward currents, primarily carried by K^+^, play an essential role as currents that restore the polarization of the cells from which a new cycle of depolarization and hyperpolarization can emerge. Consequently, the kinetics of these currents are essential in determining the overall dynamics of the oscillatory activity (Bose et al. 2014). A balance between outward and inward currents is essential for the generation of oscillatory activity: too little K^+^ current and the cell will be pushed towards, and sometimes locked in, a depolarized state; too much K^+^ current and the increased leakiness will prevent it from escaping the hyperpolarized resting state. A growing number of recent studies indicate that ionic current levels may be linked via mechanisms involving ion channel co-regulation (Bergquist et al. 2010; Linsdell and Moody 1994; MacLean et al. 2003). A consequence of this co-regulation is that, in a population of neurons, various parameters of these different currents (most notably their amplitude or conductance) are correlated with one another (Amendola et al. 2012; Anderson et al. 2016; Anirudhan and Narayanan 2015; Goaillard et al. 2009; Golowasch 2015; Khorkova and Golowasch 2007; Roffman et al. 2012; Schulz et al. 2006; Schulz et al. 2007; Srikanth and Narayanan 2015; Temporal et al. 2012; Temporal et al. 2014). Such co-regulation is likely to be involved in maintaining the balance of regenerative and outward ionic currents to regulate activity levels in oscillatory neurons.

In the stomatogastric nervous system, *I_NL_*, corresponding to the negatively sloped portion of the modulator-activated inward current *I_MI_* (Golowasch and Marder 1992b; Gray and Golowasch 2016; Swensen and Marder 2000), is a pacemaker current of the pyloric network pacemaker neurons (Bose et al. 2014; Zhao et al. 2010). This current underlies the slow oscillations observed in the presence of a variety of neuromodulators (Golowasch and Marder 1992b; Swensen and Marder 2000), even in the presence of the Na^+^ channel blocker tetrodotoxin. Although *I_MI_* is expressed by all pyloric neurons (Swensen and Marder 2000), when these neurons are synaptically isolated, modulator-induced oscillations occur only in a small subset: the three electrically coupled neurons regarded as the pacemaker neurons of the network (two Pyloric Dilators, PD, and one Anterior Burster, AB, neurons (Hooper and Marder 1987)). Why other pyloric neurons do not show pacemaker activity in the presence of modulators is unclear.

Using both theoretical and experimental methods, we test the hypothesis that the generation of slow-wave oscillations requires the correct balance of a linear pacemaker inward current and outward currents. Our hypothesis is based on the property that *I_NL_* is sufficient to emulate the pacemaker *I_MI_* (Bose et al. 2014). We show that this balance can only be produce in a subset pyloric network neurons that express the appropriate levels of high-threshold potassium currents. We further show how the induction of oscillatory activity depends on the interplay between the maximal conductance (*g_NL_*) and equilibrium potential (*E_NL_*) of *I_NL_*, and how these observations match our theoretical predictions.

## Methods

### Experimental

Experiments were performed on identified neurons from the stomatogastric ganglion (STG) of male crabs (*Cancer borealis*). The animals were obtained at local markets in Newark (NJ) and maintained in seawater tanks at 10–13°C. The entire stomatogastric nervous system, including the anterior commissural and esophageal ganglia, STG and connecting and motor nerves were dissected out as previously described (Selverston et al. 1976) and pinned down on a Sylgard-coated Petri dish, and the STG was desheathed, to allow for electrode impalement of the cell bodies. All preparations were continuously superfused with chilled (10 - 13°C) physiological *Cancer* saline: (in mM) 11 KCl, 440 NaCl, 13 CaCl_2_, 26 MgCl_2_, 11.2 Trizma base, 5.1 maleic acid, pH 7.4 –7.5.

Extracellular recordings were performed with pin electrodes placed in petroleum jelly wells built around individual nerves and recorded differentially, relative to an electrode placed outside of the well, using an A-M Systems 1700 differential amplifier (A-M Systems, Carlsberg, WA). Intracellular recordings, current injections and voltage clamp were performed with Axoclamp 2B amplifiers (Molecular Devices, Sunnydale, CA) using double impalements with 0.6 M K2SO4 + 20 mM KCl-filled borosilicate electrodes. Low resistance electrodes (15-20 MΩ) were used for current injection, and high resistance electrodes (30-40 MΩ) for voltage measurement. Individual neurons were identified by matching intracellularly recorded action potentials to action potentials on identified motor nerves that innervate known muscles (Selverston et al. 1976).

In every preparation, action potentials were blocked by bath application of 10^-7^ M tetrodotoxin (TTX; Biotium). This treatment effectively blocks all modulatory inputs, including peptidergic and cholinergic modulators that activate *I_MI_* (Golowasch and Marder 1992b; Swensen and Marder 2000), therefore decentralizing the preparation.

The dynamic clamp technique was used to activate *I*_NL_ (Fig. 1A, red) or a cut-off version of *I_NL_* that does not cross the current axis (Fig. 1A, blue) and thus corresponds to a more realistic version of the negative-slope component of *I_MI_* (Fig. 1A, gray trace). A variety of values of *I_NL_* parameters (Zhao et al. 2010) was tested. The dynamic clamp was implemented using the NetClamp software (Gotham Scientific http://gothamsci.com/NetClamp) on a 64 bit Windows 7 PC using a NI PCI-6070-E board (National Instruments).

**Figure 1.**
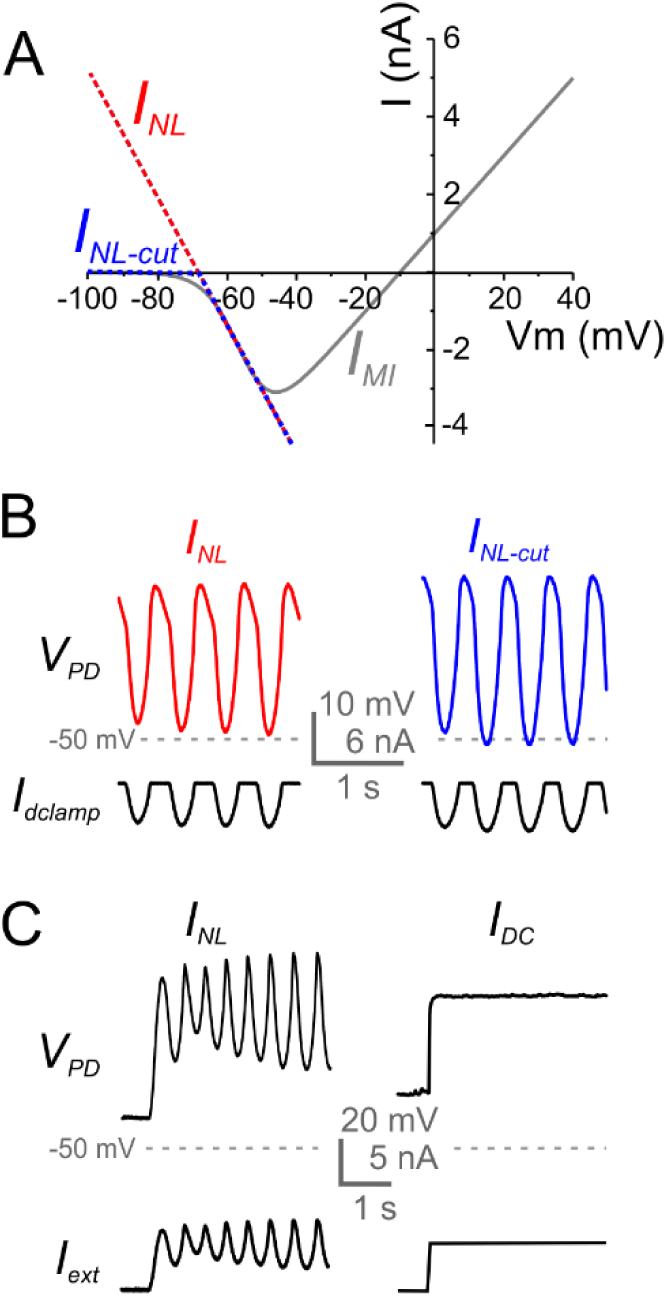
Effect of I_NL_ injection in PD neurons. **A**. The I-V curves of *I_MI_* (gray), *I_NL_* (red) and the truncated version of *I_NL_* (*I_NL-cut_*, blue). Note that both *I_NL_* and *I_NL-cut_* are good approximations to the negatively sloped portion of *I_MI_*, where

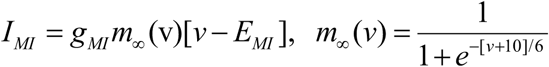 **B**. Effect of dynamic clamp injection of *I_NL_* (left) and *I_NL-cut_* (right) on the activity of PD neurons. Both traces are from the same neuron. Bottom traces show the current injected by the dynamic clamp (*I_dclamp_*). **C**. Injection of *I_NL_* (left) and comparison with injection of constant current of the same time-averaged amplitude (right) as *I_NL_*. Different preparation as in B.

Data acquisition was performed using a Digidata 1332A data acquisition board and the pClamp 10.3 software (Molecular Devices). Injections of current in dynamic clamp were performed at 10 KHz and voltage recordings at 5 KHz. The following equations were used:

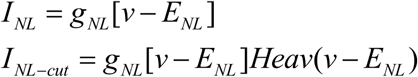

For dynamic clamp experiments involving the pyloric dilator (PD) neuron, the values used for injection were 12 values of *g_NL_* from -0.01 to -0.30 μS and the reference value of *E_NL_* was set at 2 mV below the cell’s resting potential (*V_rest_*), typically resulting in *E_NL_* = -67 to -58 mV. The value of *E_NL_* was changed by increments of ±5 mV up to ±15 mV from this reference value (a total of 7 values).

Ionic currents were measured in two-electrode voltage clamp. The high-threshold K^+^ current (*I_HTK_*) was measured with depolarizing voltage steps from a holding voltage of - 40 mV to inactivate the transient K^+^ current (*I_A_*). *I_A_* was measured with the same depolarizing voltage steps as for *I_HTK_* but from a holding voltage of -80 mV and calculated by subtraction of *I_HTK_* from these recordings (Zhao and Golowasch 2012).

Pharmacological agents were prepared immediately before use. Statistical analysis was performed with either SigmaPlot 12 (Systat) or Origin 8.5 (OribinLab) software.

### Model

The equations that describe the full model involve currents for leak (*I_L_*), negative-conductance leak with cutoff (*I_NL-cut_*) and three potassium currents, delayed rectifier (*I_Kdr_*), high-threshold (*I_HTK_*) and an A-current (*I_A_*) are:

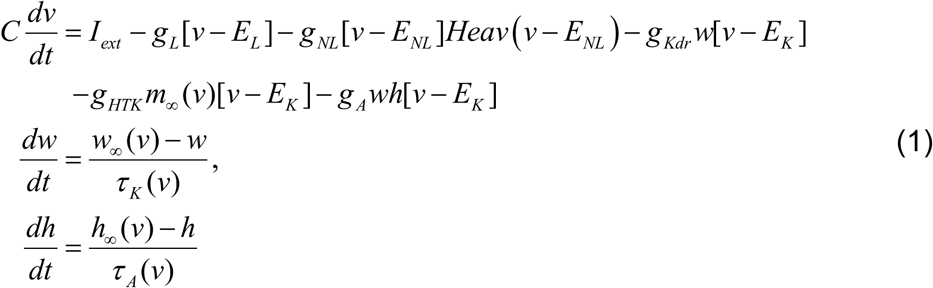

The parameter *C* is the capacitance. The variable *v* represents the membrane potential, *w* is an activation variable for potassium currents which, for convenience, is taken to be common for both *I_Kdr_* and *I_A_*, and *h* is an inactivation variable for *I_A_*. *I_HTK_* is considered to have instantaneous activation and no inactivation. The parameters *g_x_* and *E_x_* represent the maximal conductance and reversal potentials for the various currents, respectively. We use the cutoff version *I_NL_*−*cut* = *g_NL_*[*v* − *E_NL_* ]*Heav*(*v* − *E_NL_*) with a negative maximal conductance value *g_NL_* (Fig. 1A, blue). The Heaviside function *Heav*(*v* − *E_NL_*), is 0 when *v* < *E_NL_* and is equal to 1 when *v ≥ E_NL_*. This implies that for *v ≥ E_NL_, I_NL-cut_* is simply a linear current with negative conductance, while for *v* < *E_NL_*, *I_NL-cut_* = 0 (Fig. 1A, blue). The terms *w*_∞_(*v*) and *m*_∞_(*v*) are the steady-state activation functions for the two potassium currents and *h*_∞_(*v*) is the steady state inactivation function of *I_A_*. They are described by equations:

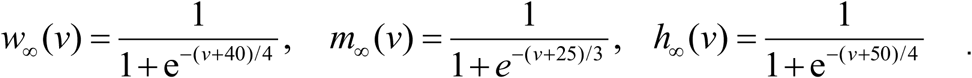

The associated time constants are given by

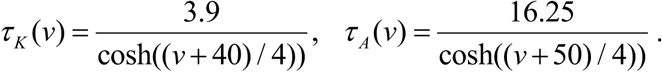

Parameter values that vary across simulations are provided in the Results, while those that were fixed are given here: *C* = 1 μF/cm^2^, *g_L_* = 0.00325 μS, *E_L_* = -60 mV, *g_Kdr_* = 0.0325 μS, *E_K_* = -80 mV and *I_ext_* = 0.065 nA. These parameters were chosen so that the period of oscillations in the simulations was on the order of those found in experiments. Simulations and bifurcation diagrams were constructed using XPPAUT (Ermentrout 2002).

In prior work (Bose et al. 2014), we analyzed the case of *I_HTK_* = *I_A_*= 0 and showed that non-zero *I_NL_* and *I_Kdr_* currents alone can produce oscillations. In order to now isolate the effect of *I_HTK_* with regard to pacemaker properties, for much of the analysis we continue to keep *I_A_*=0. With *I_A_*=0, the variable *h* is redundant and the set of equations (1) reduces to a two-dimensional system that will be analyzed using phase plane methods. We will show that a non-zero value of *I_A_* does not affect the existence of oscillations, but that if it is large enough it can introduce a stable, sub-threshold fixed point.

## Results

### The pacemaker current operates over restricted parameter ranges

Our first goal here is to characterize some of the conditions that *I_MI_* needs to satisfy to operate as a pacemaker current as predicted by our previous theoretical work (Bose et al. 2014). Oscillatory activity can be induced in the PD neuron by injecting the negative-leak conductance current *I_NL_* (Fig. 1B, C), which is a linearized version of the pacemaker current *I_MI_* (Fig. 1A). The effect of *I_NL_* only depends on the region of its I-V curve where the current is negative. Therefore, injection of the same current which is set to 0 below *E_NL_* (the cut-off version *I_NL-cut_*) produces almost identical oscillations (Fig. 1B; see Methods). Since the results for oscillatory activity are nearly identical, henceforth we use *I_NL_*.

The effect of *I_NL_* in producing oscillations in the PD neuron could not be mimicked by injecting a depolarizing DC current (Fig. 1C right) equal to the time-averaged current measured during the dynamic clamp injection of *I_NL_* (Fig. 1C left), demonstrating that the effects of *I_NL_* are not simply a consequence of a depolarization of the cell by *I_NL_*, but its role as a voltage-dependent current.

In our previous study, we predicted that oscillations produced by *I_NL_* would occur in a restricted range of *g_NL_* (Bose et al. 2014). To explore the effect of the *I_NL_* parameters on the oscillatory activity, we changed *g_NL_* over 12 values from -0.01 to -0.30 μS and *E_NL_* over a range of ±15 mV in steps of ±5 mV from the initial reference value (for a range of 12 x 7 = 84 runs; see Methods). We found that there was a double Gaussian distribution (R^2^=0.9185) of *g_NL_*-*E_NL_* values over which *I_NL_* was effective in eliciting oscillatory activity (Fig. 2A).

An important point that these results illustrated was that oscillatory activity in the pyloric pacemaker cells does not require TTX-sensitive Na^+^ currents to be produced. We observed a relative independence of cycle period on *g_NL_* (Fig. 2B), but a decreasing amplitude of oscillations as *g_NL_* became larger in absolute value (Fig. 2C). On the other hand, period was an increasing function of *E_NL_* (Fig. 2D) and the oscillation amplitude had an inverted U-shape relationship with *E_NL_* (Fig. 2E).

**Figure 2.**
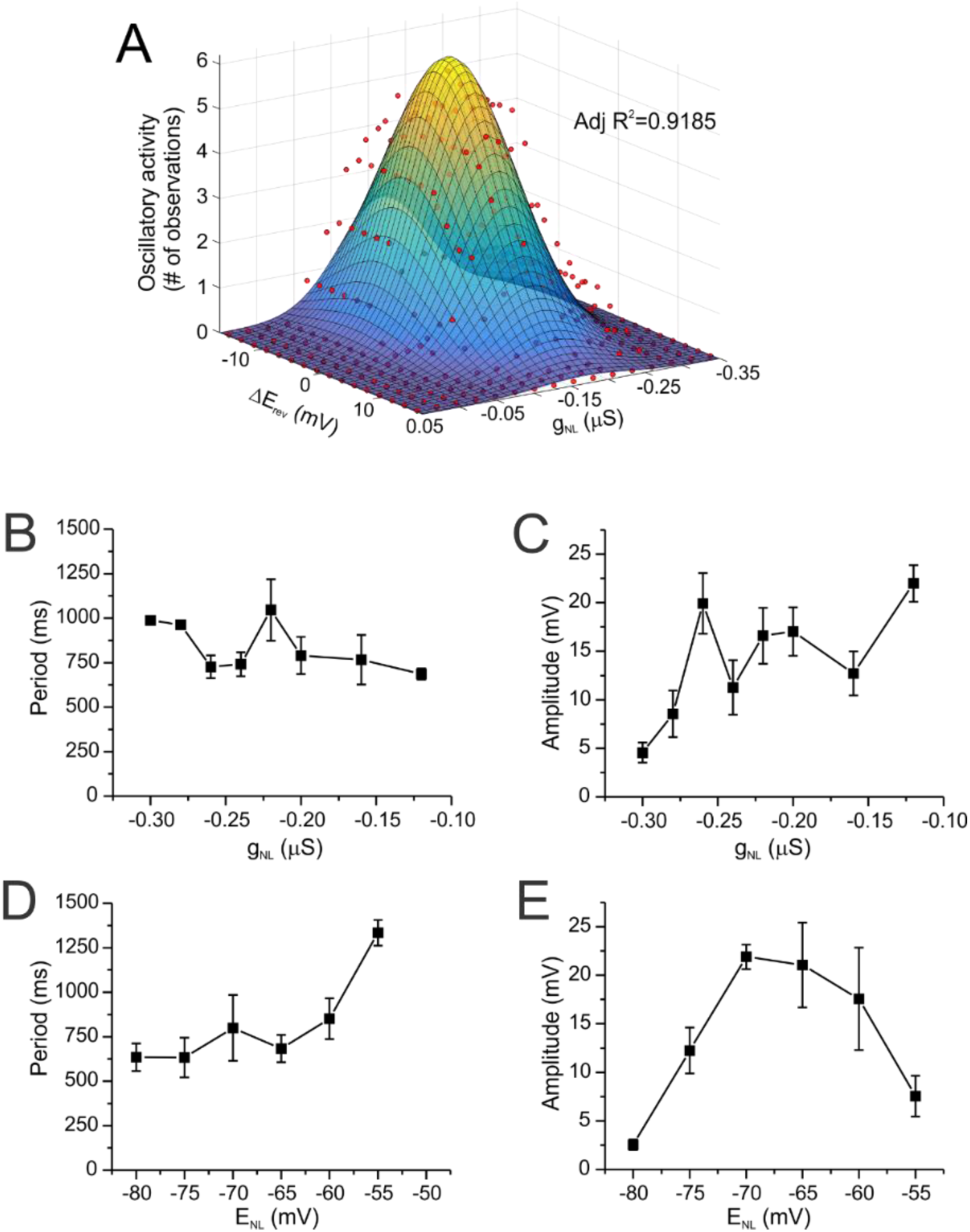
Effect of I_NL_ parameters on PD neuron oscillations. Identified PD neurons (N=7) were injected with dynamic clamp *I_NL_* and parameters *g_NL_* and *E_rev_* were modified over a broad range of values. **A**. The presence or absence of oscillations (# of oscillatory preparations out of 7) was graphed as a function of the change in *E_rev_* relative to each cell’s resting voltage (*ΔErev* = 0 = *V_rest_* – 2 mV), and the value of the negative conductance injected (*g_NL_*). Red symbols are the experimental data. The smooth surface is a Gaussian surface fit to the experimental data. Adjusted R^2^ = 0.9185. **B**. Average period vs *g_NL_*. **C**. Average amplitude vs *g_NL_*. **D**. Average period vs *E_NL_*. **E**. Average amplitude vs *E_NL_*. All data shown in A-E are from the same set of cells. Error bars are SEM.

### Pacemaker cells balance inward and outward currents to produce oscillatory activity

#### The effect of I_HTK_

As we have seen, *I_NL_* injected into neurons of the pacemaker group (i.e. PD neurons) consistently induces oscillatory activity (Fig. 2, Fig. 3, top left), albeit within relatively narrow ranges of *g_NL_* and *E_NL_* (Fig. 2). We examined whether follower neurons in the network were equally capable of generating oscillatory activity.

**Figure 3.**
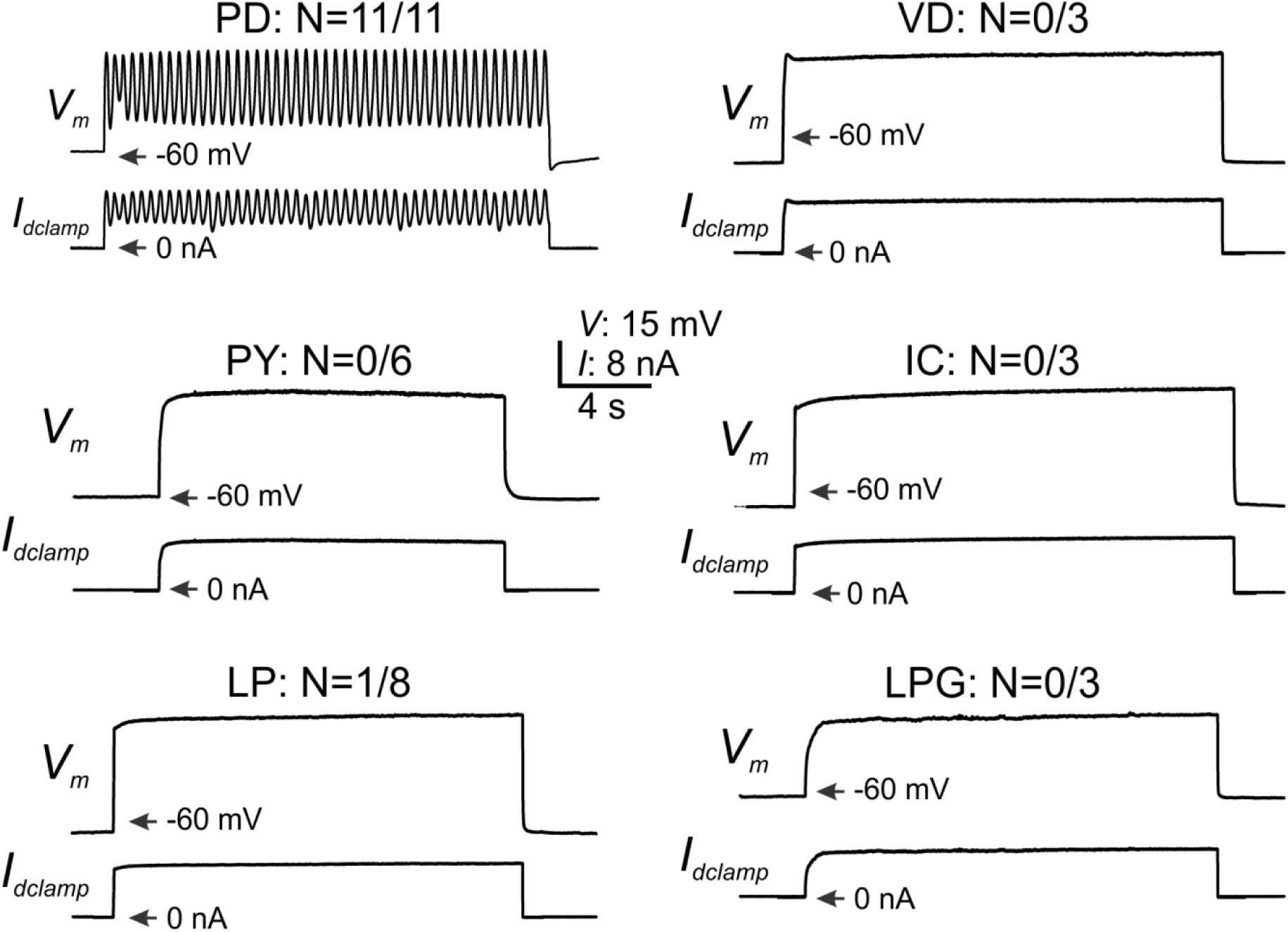
D*ynamic clamp I_NL_ injection in pyloric network neurons cannot induce oscillatory activity in follower cells*. Identified neurons with characteristic activity response to *I_NL_* injection (*g_NL_* = 0.16 μS, *E_NL_* = *V_rest_* – 2 mV) of the cells listed. PD neurons responded with oscillatory activity in 100% of the preparations tested. The numbers next to the cell type name indicate how many of the total cells tested showed any oscillatory activity. All cells were tested with the same combinations of *g_NL_* and *ΔE_rev_* as the PD neurons shown in Figure 2A. Top traces in each panel is the membrane potential; bottom trace is the dynamic clamp current injected.

We observed that none of the follower cells in the pyloric network (ventral dilator VD, pyloric constrictor PY, inferior cardiac IC, lateral pyloric LP and lateral posterior gastric LPG neurons) were capable of generating consistent oscillatory activity independently of the combination of *g_NL_* and *E_NL_* used. Individual examples shown in Figure 3 were obtained by injection of *g_NL_* = -0.16 μS and *E_NL_* = *V_rest_* – 2 mV for each cell type. However, each cell was further tested with the same combination of *g_NL_* and *ΔE_rev_* as the PD cells shown in Figure 2A. We found that only one out of eight LP neurons tested expressed any oscillatory activity, and none of the other follower cells tested could express such activity for any of the *g_NL_* and *E_NL_* combinations.

To understand what prevents follower cells from expressing oscillatory activity, we examined the levels of K^+^ currents expressed in two of the follower cells (LP and VD neurons) and compared that to the current levels expressed by the pacemaker PD neurons. Our hypothesis throughout was that large outward currents could be responsible for preventing oscillatory activity. Figure 4 shows the comparison of two outward currents, *I_HTK_* and *I_A_* (see definitions in Methods), with *I_HTK_* further divided into peak (*I_HTK_* peak) and steady state (*I_HTK_* SS) in two cell types the PD and LP neurons.

PD neurons expressed a significantly smaller *I_HTK_* than LP neurons, when this current was compared at its peak (two-way ANOVA, P = 0.029, Fig. 4B), as well as at steady state (P < 0.001, Fig. 4C). Interestingly, the activation parameters as well as the maximal conductance of the peak of *I_HTK_* are not significantly different between PD and LP neurons (Fig. 4E; PD, Table 1). However, the differences, especially in *g_max_HTK_* and *V_1/2_HTK_*, while independently not significantly different between these cells, were sufficient to make the currents different between them.

**Figure 4.**
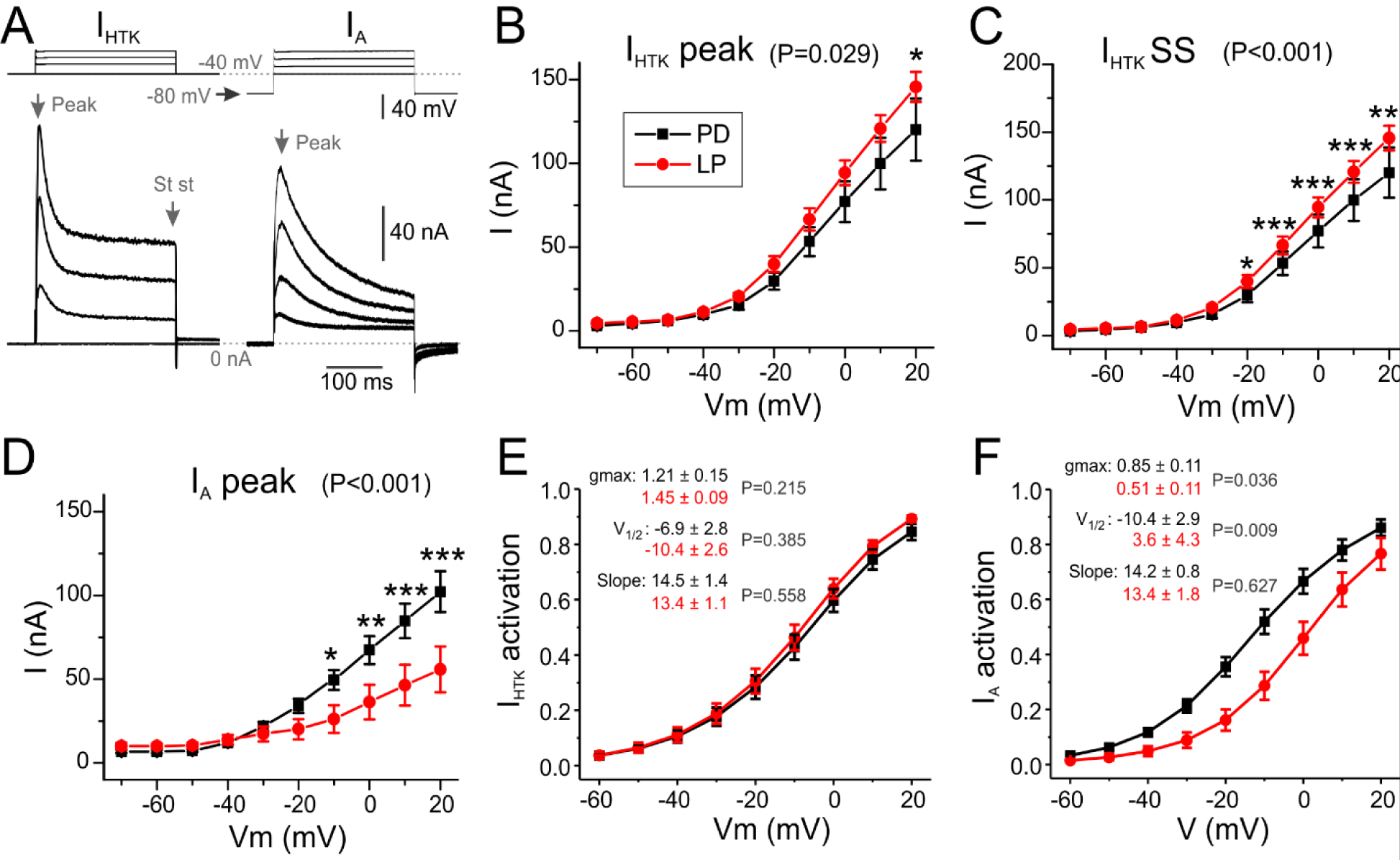
Potassium current levels in two key pyloric network neurons. **A**. Sample traces of *I_HTK_* (left) and *I_A_* (right). Grey vertical arrows point to the times at which *I_HTK_ peak*, *I_HTK_ SS* and *I_A_* are measured. **B**. I-V curves of the peak of the high threshold current *I_HTK_*. The curves are significantly different (Two-way ANOVA, P < 0.029). **C**. Steady state I-V curves of *I_HTK_*. The curves are significantly different (Two-way ANOVA, P < 0.001). Asterisks indicate the source of the difference from two-way ANOVA analysis and Tukey pairwise comparisons (* P < 0.05; ** P < 0.01; *** P < 0.001). **D**. I-V curves of the peak of the transient K^+^ current *I_A_*. The curves are significantly different (Two-way ANOVA, P < 0.001).). Data for PD neurons in black and red for LP neurons. **E**. Activation curves for *I_HTK_* peak and the shown parameters are calculated as in D. **F**. Average of the voltage-dependent activation curves of *I_A_* derived from the I-V curves of each cell used in A. The I-V curves were fitted with the equation. 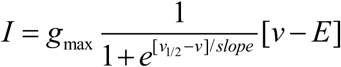. *E* was fixed at -80 mV and the other three parameters were determined from the fit (*V_1/2_* and *slope* in mV, *g_max_* in μS. P values are for Students t-test analysis for each parameter between the two cells. All error bars are SEM.

**Table 1:** *I_HTK_* and *I_A_* parameters in PD and LP neurons. Parameters were obtained from fits of a sigmoidal function to conductances calculated from *I_HTK_* and *I_A_* measurements and a driving force = (*V_m_* - *E_rev_*), with *E_rev_* = -80 mV.

|  | Mean |  | SD |  | t (d.f.) | P |
| --- | --- | --- | --- | --- | --- | --- |
|  | PD | LP | PD | LP |  |  |
| $g_{max\_HTK}$ (μS) | 1.21 | 1.45 | 0.61 | 0.33 | -1.270 (28) | 0.215 |
| $V_{1/2\_HTK}$ (mV) | -6.9 | -10.4 | 11.6 | 9.3 | 0.883 (28) | 0.385 |
| $V_{slope\_HTK}$ (mV) | 14.5 | 13.4 | 5.8 | 3.8 | 0.593 (28) | 0.558 |
| $g_{max\_A}$ (μS) | 0.85 | 0.51 | 0.44 | 0.39 | 2.205 (28) | 0.036 |
| $V_{1/2\_A}$ (mV) | -10.4 | 3.6 | 11.8 | 15.7 | -2.829 (28) | 0.009 |
| $V_{slope\_A}$ (mV) | 14.2 | 13.5 | 3.4 | 6.5 | 0.491 (28) | 0.627 |

In contrast to *I_HTK_*, PD neurons had a significant larger *I_A_* than LP neurons (two-way ANOVA, P = 0.001, Fig. 4D) in part derived from a significantly more hyperpolarized activation curve (*V_1/2_A_*, Fig. 4F, Table 1) and a significantly higher maximum conductance (*g_max_A_*,Table 1). The *V_slope_* of *I_A_* did not differ significantly between the two cells (Table 1).

#### The functional significance of K^+^ current amplitude differences

In order to test if the differences in K^+^ current levels reported above are functionally related to the inability of follower cells to generate oscillatory activity when injected with *I_NL_*, we reduced *I_HTK_* with different concentrations of TEA (Golowasch and Marder 1992a) in the LP neuron. A significant effect of TEA on the *I_HTK_* I-V curve is observed at all voltages (two-way RM ANOVA, P = 0.008, Fig. 5A). In fact, a highly significant effect of TEA concentrations was observed, with a maximum inhibition level of ∼80% and a half-maximal effect at 1.6 mM (Fig. 5B). We tested whether reduced *I_HTK_* conditions were more permissive for producing oscillations by injecting *I_NL_* (*g_NL_* = -0.16 μS) in the presence of 8 mM TEA. We observed that, under these conditions, oscillatory activity could be consistently elicited in the LP neuron (Fig. 5C; N = 4) comparable to those generated by PD neurons (compare with Fig. 1B, C). As with PD neurons (see Fig. 1C), DC injections could not trigger oscillatory activity in LP neurons (Fig. 5D).

**Figure 5.**
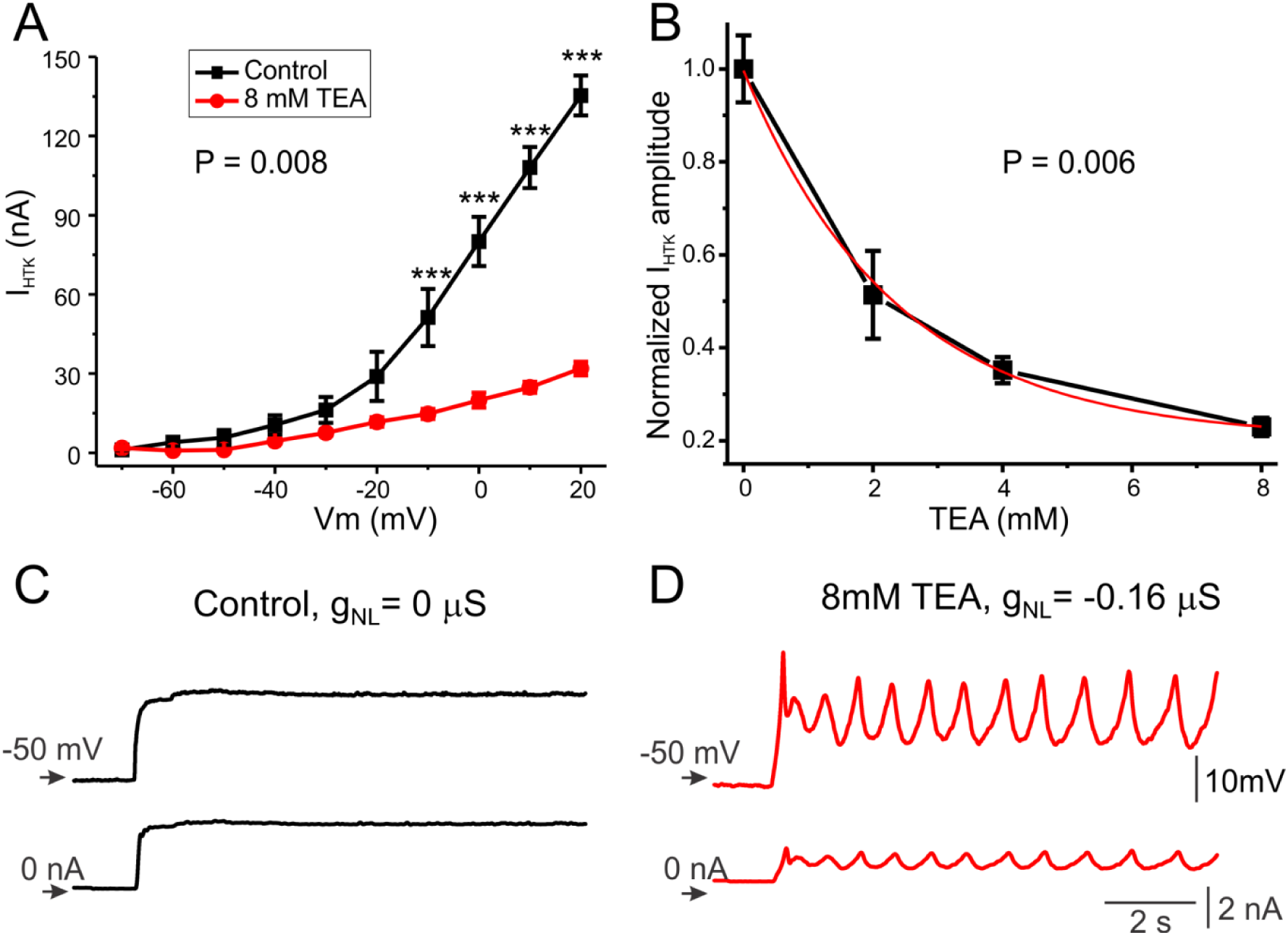
High I_HTK_ levels are responsible for the inability of LP neurons to generate oscillatory activity. **A.** I-V curves of the peak *I_HTK_* in LP neurons under control (0 mM TEA, black) and 8 mM TEA (red). A significant decreases is observed overall (Two-way RM ANOVA, N = 3, P = 0.008) with asterisks indicating the source of the difference from Tukey pairwise comparisons (*** P < 0.001). **B**. Dose-dependent sensitivity of peak *I_HTK_* to TEA. *I_HTK_ peak* was normalized to the current measured in control (0 mM TEA). Average ± SEM is plotted. The overall effect was statistically significant (One-way RM ANOVA, N = 3, P = 0.006). The red trace is an exponential fit with a saturation level at ∼20%, and an ID_50_ = 1.6 mM. **C**. Effect of dynamic clamp injection of *I_NL_* (*g_NL_* = -0.16 μS; *E_NL_* = -52 mV). Top trace is membrane potential, bottom trace is current injected by the dynamic clamp circuit. **D**. Effect of injection of DC equal to the time-averaged current during *I_NL_* injection shown in C.

#### The effect of I_A_ current amplitude differences

Because *I_A_* also shows significantly different levels in PD neurons compared to the LP neuron, albeit with higher levels in the PD neuron (Fig. 4A), we tested the effect of modifying *I_A_* using the blocker 4-AP in PD neurons. We expected that, if *I_A_* is involved in the regulation of PD neurons to generate oscillatory activity, reducing *I_A_* levels in the PD neuron would affect its oscillatory properties. Figure 6 shows that 1 mM 4-AP significantly affects *I_A_* overall (two-way ANOVA, P = 0.012 for 4-AP treatment), reducing it to levels similar to those measured in LP neurons, i.e. ∼50% at all voltages tested (see Fig. 4A). Yet, applying 1 mM 4-AP had virtually no effect on the ability of these cells to oscillate in response to *I_NL_* injection (compare Fig. 6B control with similar injection in the presence of 1 mM 4-AP in Fig. 6C).

**Figure 6.**
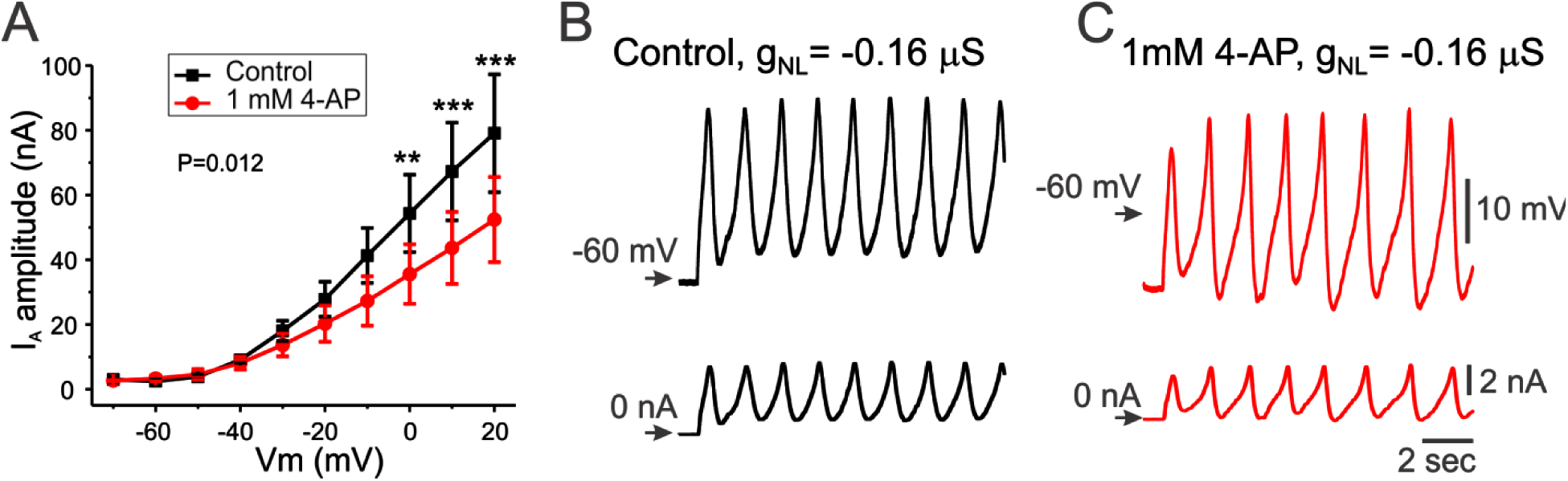
I_A_ does not appear to be involved in the ability of PD neurons to generate oscillatory activity. **A.** I-V curves of the peak *I_A_* in PD neurons under control (0 mM 4-AP, black) and 1 mM 4-AP (red). A significant decrease is observed overall (Two-way RM-ANOVA, N = 3, P = 0.012) with asterisks indicating the source of the difference using Tukey pairwise comparisons (** P < 0.01, *** P < 0.001). **B**. Effect of dynamic clamp injection of *I_NL_* in the absence of 4-AP (*g_NL_* = -0.16 μS; *E_NL_* = -72 mV). Top trace is membrane potential, bottom trace is current injected by the dynamic clamp circuit. **C**. Effect of dynamic clamp injection of *I_NL_* in the presence of 1 mM 4-AP (*g_NL_* = -0.16 μS; *E_NL_* = -72 mV).

Similar results were observed when comparing the PD and VD neurons, the latter another follower cell in the pyloric network. *I_HTK_* in PD neurons was found to be significantly lower in amplitude across its voltage activation range than in VD neurons (two-way ANOVA, P = 0.001, Fig. 7A), whereas, unlike the comparison of PD and LP neurons, there was no significant difference in *I_A_* between these two cells (two-way ANOVA, P = 0.787, Fig. 7B).

**Table 2:**
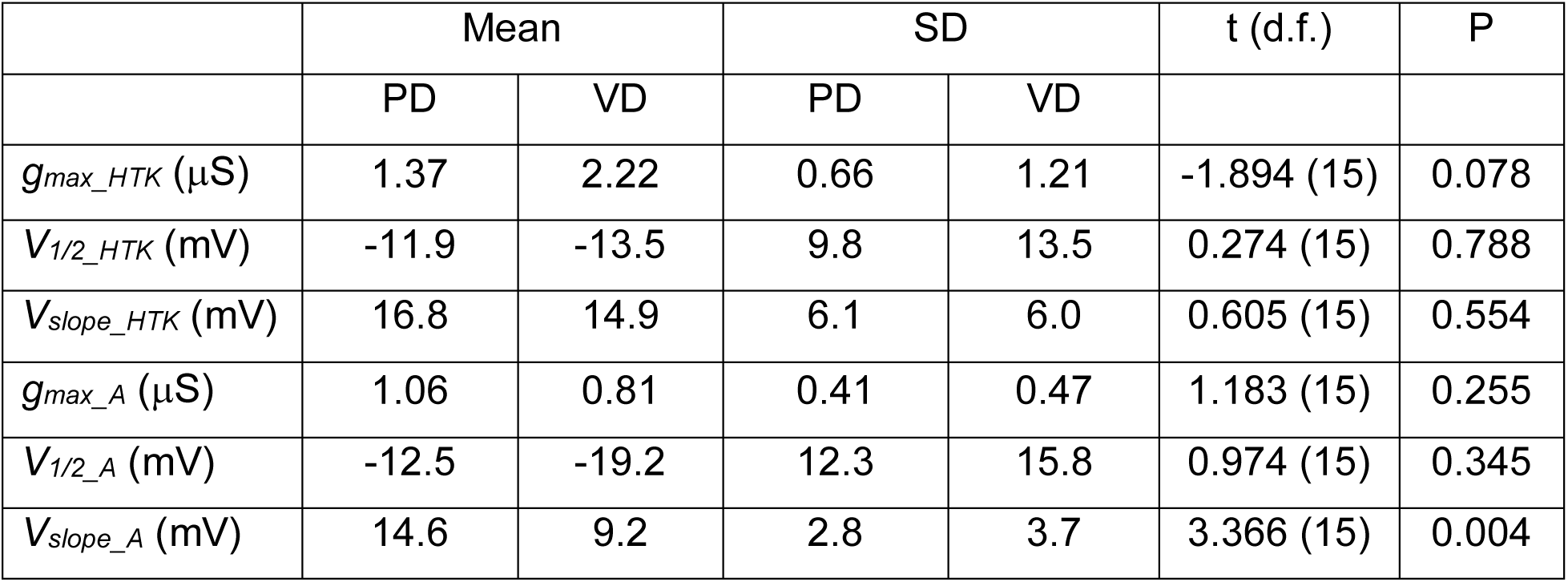
*I_HTK_* and *I_A_* parameters in PD and VD neurons. Parameters were obtained from fits of a sigmoidal function to conductances calculated from *I_HTK_* and *I_A_* measurements and a driving force = (*V_m_* - *E_rev_*), with *E_rev_* = -80 mV.

|  | Mean |  | SD |  | t (d.f.) | P |
| --- | --- | --- | --- | --- | --- | --- |
|  | PD | VD | PD | VD |  |  |
| $g_{max\_HTK}$ (μS) | 1.37 | 2.22 | 0.66 | 1.21 | -1.894 (15) | 0.078 |
| $V_{1/2\_HTK}$ (mV) | -11.9 | -13.5 | 9.8 | 13.5 | 0.274 (15) | 0.788 |
| $V_{slope\_HTK}$ (mV) | 16.8 | 14.9 | 6.1 | 6.0 | 0.605 (15) | 0.554 |
| $g_{max\_A}$ (μS) | 1.06 | 0.81 | 0.41 | 0.47 | 1.183 (15) | 0.255 |
| $V_{1/2\_A}$ (mV) | -12.5 | -19.2 | 12.3 | 15.8 | 0.974 (15) | 0.345 |
| $V_{slope\_A}$ (mV) | 14.6 | 9.2 | 2.8 | 3.7 | 3.366 (15) | 0.004 |

**Figure 7.**
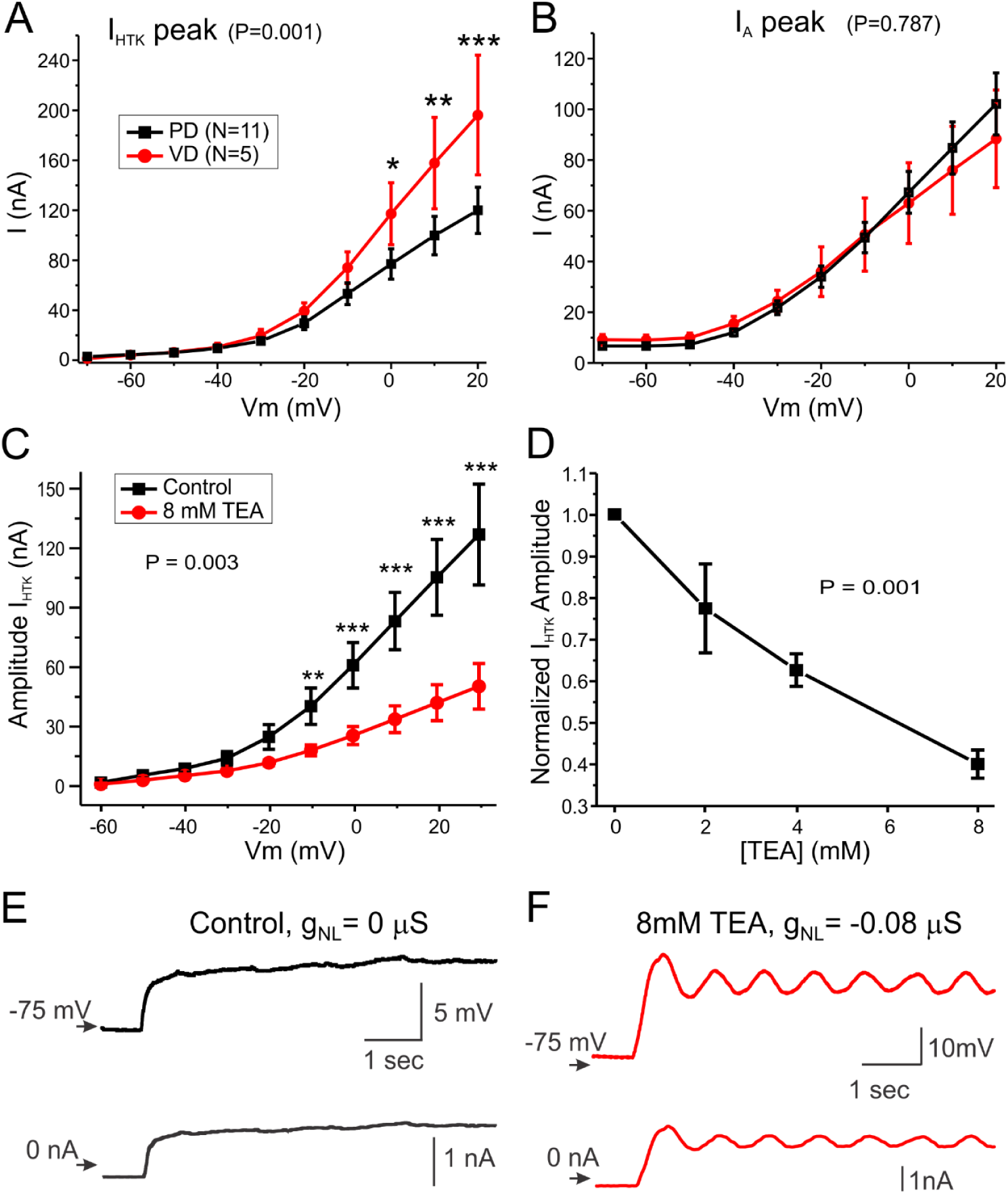
High I_HTK_ levels are responsible for the inability of VD neurons to generate oscillatory activity. **A.** I-V curves show that the peak *I_HTK_* in VD neurons is almost twice as large as in PD neurons (Two-way ANOVA, P = 0.001) over most of its activation range. **B**. I-V curves of *I_A_* show no significant difference between PD and VD neurons (Two-way ANOVA, P = 0.787). **C.** I-V curves of the peak *I_HTK_* in VD neurons under control (0 mM TEA, black) and 8 mM TEA (red). A significant decrease is observed overall (Two-way RM ANOVA, N = 4, P = 0.003) with asterisks indicating the source of the difference from a Tukey pairwise comparisons. * P < 0.05** P < 0.01; *** P < 0.001). **D**. Sensitivity of *I_HTK-peak_* to TEA. *I_HTK-peak_* was normalized to the current measured in control (0 mM TEA). Average ± SEM is plotted. The overall reduction of *I_HTK-peak_* by TEA was significant (One-way RM ANOVA, N = 4, P = 0.001). **E, F**. Effect of dynamic clamp injection of *I_NL_* (*g_NL_* = -0.08 μS; *E_NL_* = -70 mV) in normal saline (**E**) and in the presence of 8 mM TEA (**F**). Top trace is membrane potential, bottom trace is current injected by the dynamic clamp circuit.

Bath application of 8 mM TEA reduced the amplitude of *I_HTK_* significantly (Fig. 7C, N=4, two-way RM-ANOVA, P = 0.003). The effect of TEA was dose-dependent (one-way RM-ANOVA, P < 0.001, Fig. 7D). When *I_NL_* was injected (*g_NL_* = -0.16 μS) in control medium (0 mM TEA), VD neurons such as the one shown in Figure 7E depolarized but never generated oscillations (see also Fig. 3). However, in the presence of 8 mM TEA, the same *I_NL_* injection elicited oscillations (Fig. 7F) comparable to those produced by *I_NL_* in PD neurons in normal TTX saline (Fig. 1B, C) and LP neurons in TEA (Fig. 5D).

### Modeling description of experimental observations

We performed simulations using equation (1) to determine whether a theoretical model provides a framework for understanding our experimental findings. The main question we investigated was whether differences in various K^+^ currents could cause a model neuron to either oscillate or not. Additionally, we checked to see what computational predictions our model made with regards to period and amplitude as functions of parameters associated with *I_NL_*.

To guide the simulations, we first discuss the phase space structure of the model. Recall that we are keeping *g_Kdr_* fixed throughout at 0.0325 μS, and *g_L_* at 0.00325 μS. This is to ensure oscillations in the absence of additional K^+^ currents when *I_NL_* is added.

Now we examine the role of other K^+^ currents known to be expressed in most neurons, and in pyloric neurons in particular. When *I_A_* = 0, the model equations involve only the *v* and *w* variables, allowing for phase plane analysis. The nullclines of equation (1) are obtained by plotting the set of points that satisfy *v*’=0 and *w*’=0, respectively. The *v*- nullcline is decreasing for *v* < *E_NL_*. If |*g_NL_*| is sufficiently large relative to *g_L_*, then the nullcline can increase for a range of values *v ≥ E_NL_*. The *w*-nullcline is a monotone increasing sigmoidal function. If it intersects the *v*-nullcline along its decreasing portion, then a stable fixed point ensues and oscillations are not possible. If the intersection of the two nullclines occurs for *v ≥ E_NL_* then oscillations may be possible and depends on the slope of each at the point of intersection. We will show that, as *g_HTK_* is increased, oscillatory behavior is destroyed.

Figure 8A1 shows phase plane from equation (1) when *g_NL_* = -0.0195 μS, *g_HTK_* = 0.013 μS and *g_A_* = 0 μS. For this choice of parameters, the *w*-nullcline intersects the *v*- nullcline along a portion where the latter is increasing with a sufficiently large slope at the point of intersection. This produces an unstable fixed point. A periodic solution surrounds this unstable fixed point because, as *v* becomes too hyperpolarized, *I_NL_* current drives the voltage away from *E_NL_*. At larger values of *v*, *I_Kdr_* activates and restricts the voltage from increasing too much. This result is consistent with our prior work (Bose et al. 2014) in which *I_HTK_* was not present. Thus, the addition of this small amount of *HTK* conductance does not play a role in the generation of oscillations, but it does limit the amplitude at both higher and lower values of *v*. Indeed, the largest amplitude oscillation under these conditions exists when *g_HTK_* = 0 (Fig. 8B).

**Figure 8.**
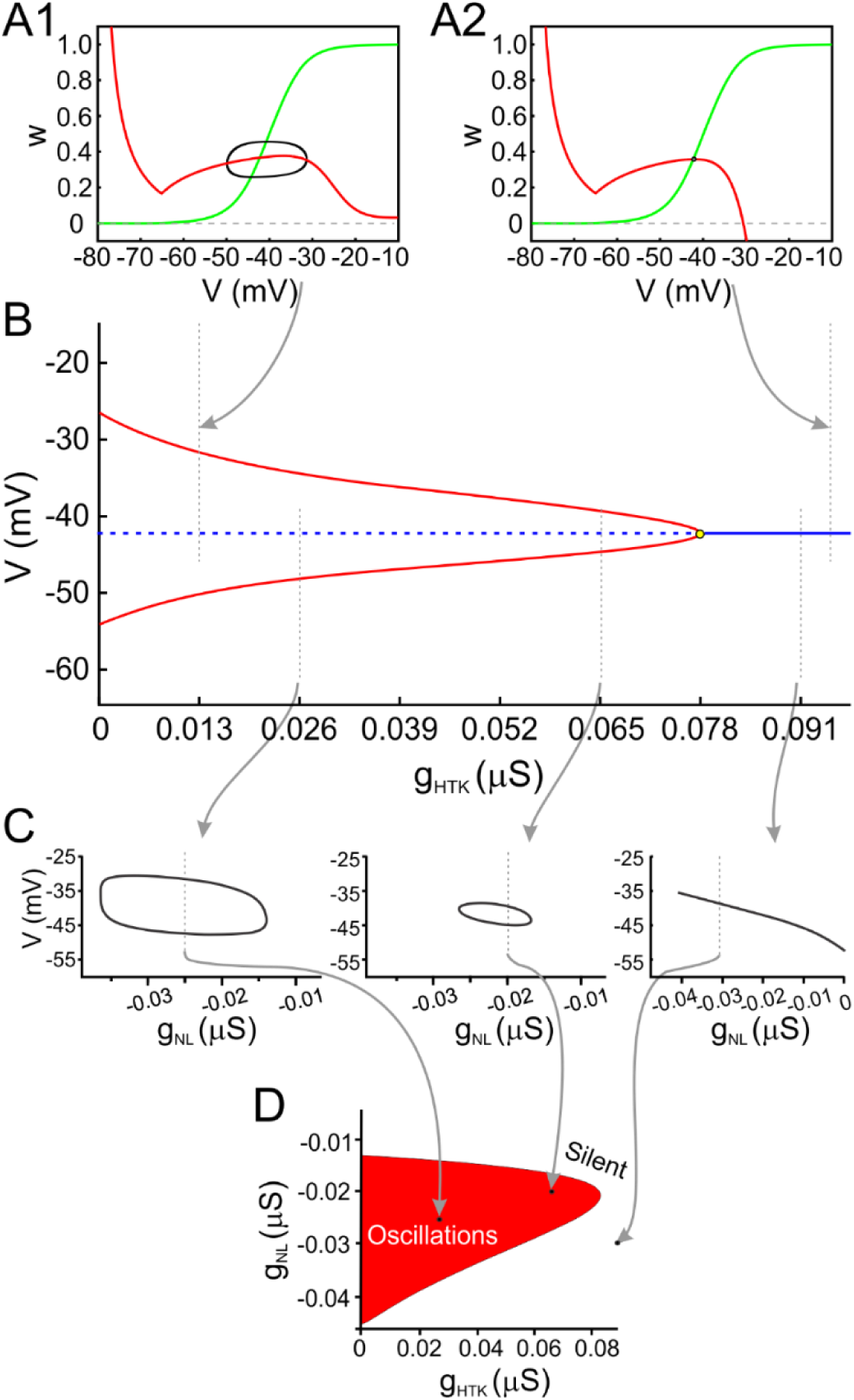
The effect of I_HTK_ on the existence of oscillations. **A**. Panel A1 shows the v-w phase plane for low values of *g_HTK_* in which an oscillatory solution exists. For larger values of *g_HTK_* the oscillatory solution fails to exist, as shown in panel A2, because the stronger *I_HTK_* stabilizes the fixed point at the intersection of the two nullclines. **B**. Bifurcation diagram showing how the existence and amplitude of oscillatory solutions depends on *g_HTK_*. Arrows from panels A1 and A2 show the values of *g_HTK_* used to produce those simulations. Oscillatory solutions are lost through a Hopf Bifurcation as *g_HTK_* increases. **C**. Bifurcation diagrams showing behavior of solutions at gixed values of *g_HTK_* while *g_NL_* is varied. The range of *g_NL_* values for which oscillations exist decreases at *g_HTK_* increases (compare left and middle diagrams). If *g_HTK_* is too large (rightmost diagram), no amount of *g_NL_* can restore oscillations. **D**. Bifurcation diagram summarizing behavior in the *g_HTK_*- *g_NL_* parameter space. Oscillations are possible for parameter values in the red region bounded between the parabola-like shape and the vertical axis and are not possible outside of that region.

We then increased the value of *g_HTK_* to 0.0975 μS. Figure 8A2 shows the ensuing phase plane. Note that the only change is that the *v*-nullcline now has a much steeper and pronounced right branch along which it is decreasing. Moreover, the slope of the *v*- nullcline at the intersection with the *w*-nullcline has decreased enough to stabilize that fixed point. This change is indicative of a Hopf bifurcation, which in fact occurs as *g_HTK_* is increased. Figure 8B shows that as *g_HTK_* is increased from lower values, the periodic solution decreases in amplitude (upper and lower bounds of oscillations shown by solid red curves). Moreover, the unstable fixed point (dashed blue curve) gains stability through a Hopf bifurcation near *g_HTK_* =0.078 μS and oscillations cease to exist. The reason for this loss of excitability is that *I_HTK_* is too strong and destroys the balance between *I_Kdr_* and *I_NL-cut_* that could produce oscillations. In short, too much *I_HTK_* is incompatible with the production of oscillations. This finding is consistent with our experimental results shown in Figs. 5 and 6.

We next varied *g_NL_* to further understand the relationship between these currents. Figure 8C shows bifurcation diagrams for three different fixed values of *g_HTK_*, indicated by arrows from Fig. 8B, as we vary *g_NL_*. The rightmost panel shows that if *g_HTK_* is too large, then no amount of *I_NL-cut_* can produce oscillations. Indeed for all values of *g_NL_*, the stable fixed point remains. For smaller values of *g_HTK_* (middle and left panels), oscillations are present over an interval of *g_NL_* values. This interval is a decreasing function of *g_HTK_* as summarized in Fig. 8D. There we show a two-parameter bifurcation diagram where the curve is the boundary between oscillatory and non-oscillatory behavior. For any choice of parameters that lies within the red shaded parabola-like region, oscillations are possible. For choices outside of this region, oscillations are not possible. The diagram clearly shows that the interval of *g_NL_* values over which oscillations occurs shrinks as *g_HTK_* is increased.

Our experimental findings, as summarized in Fig. 7, suggest that *I_A_* plays no role in generating oscillations, but may affect the properties of the oscillatory solution. Therefore, we investigated what role, if any, *I_A_* may play in the generation of oscillations. With a non-zero value of *I_A_*, equation (1) becomes three-dimensional and direct phase plane analysis is not possible. Thus, we now project the solution trajectory onto the *v-w* phase plane to study how the shape of the respective nullclines varies as a function of parameters associated with *I_A_*. For example, Figure 9A1, shows the case where *g_A_*= 0.0325 μS. Note that the projected nullclines look very much the same as those in Figure 8A1. As we varied the value of either *gA* or the half-activation voltage of *h*_∞_(*v*) we found little change in the shape of the projected nullclines. Thus, we concluded that the addition of *I_A_* plays no role in whether a periodic solution exists, consistent with results of the prior sub-section. However, an interesting effect of a large *I_A_* current was found to occur. A very strong *A*-current can induce bistability between a periodic solution and a stable fixed point. Figure 9A2 shows the case of *g_A_* = 0.195 μS. For this parameter value, a stable periodic solution co-exists with a stable fixed point. The figure shows an example of a trajectory that starts at the stable fixed point where *h* = 0.45 (lower *v-* nullcline). Leaving the initial values of *v* and *w* unchanged, we switched to the initial condition *h* = 0. This has the effect of assigning a new initial condition in the full three-dimensional v-w-h phase space that is not a fixed point (though the location looks unchanged in the projection onto the v-w phase plane). The ensuing trajectory leaves that location and is seen to converge to the stable periodic orbit. The associated bifurcation diagram in Figure 9B shows that the stable fixed point loses stability through a subcritical Hopf bifurcation as *g_A_* is decreased. The red, roughly horizontal curves depict the upper and lower limits of the periodic solution.

**Figure 9.**
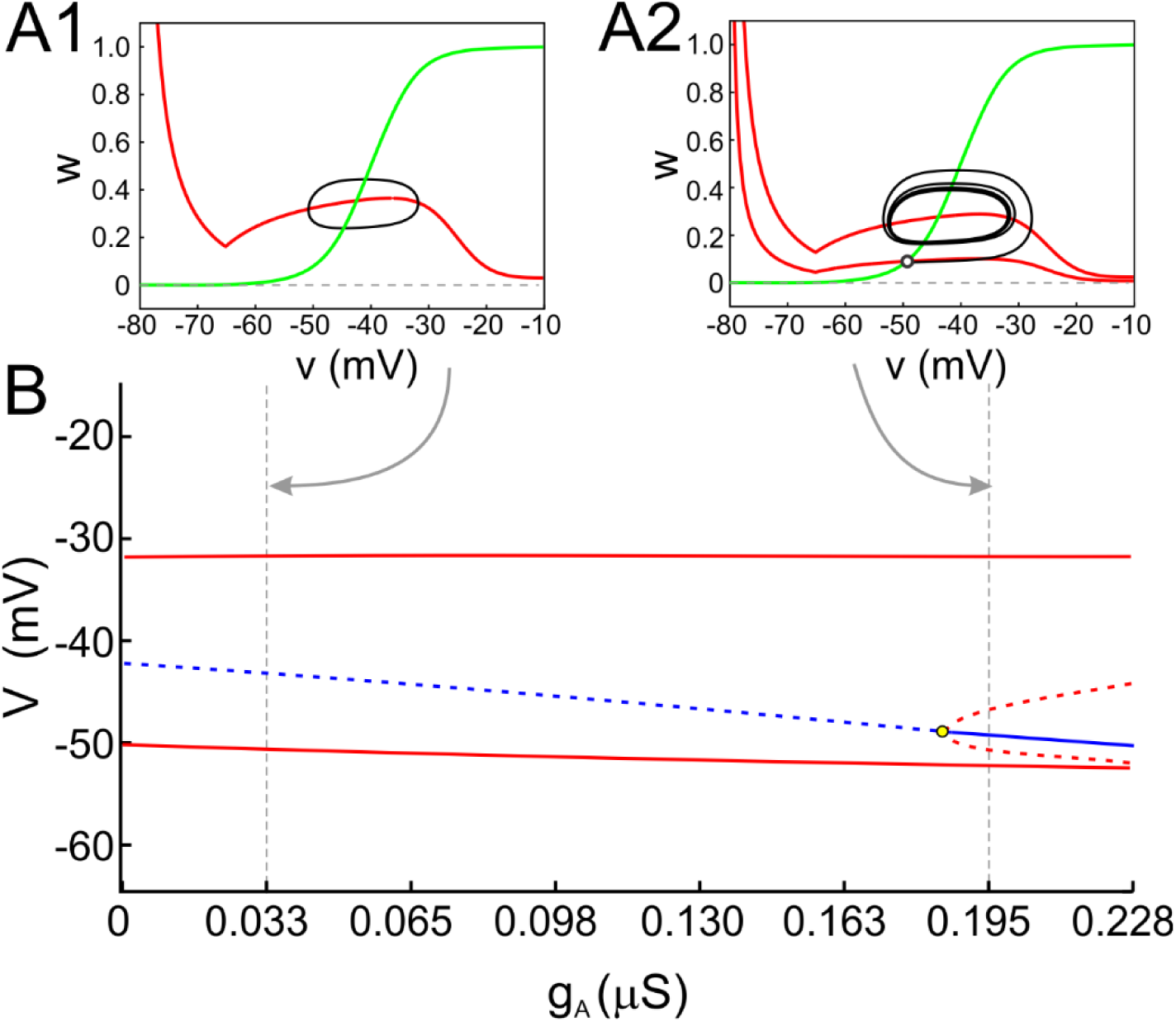
The effect of I_A_ on the existence of oscillations. **A1**.The projection onto the *v-w* phase plane for *g_A_*=0.033 μS. No qualitative difference is found from the *g_A_* = 0 μS case (shown in Fig. 8A1). The value of the third variable *h* (not shown) oscillates, but remains small. The *v*–nullcline shown is for the largest value of *h* along the periodic solution. The intersection of the projection of the *v* and *w*-nullclines is not a fixed point. **A2**. The projection onto the *v-w* phase plane at *g_A_*=0.195 μS for which bistability exists between the periodic solution and a stable fixed point. The lower of the two *v*-nullclines occurs for *h* = 0.45 at the value corresponding to a stable fixed point (open circle). The upper of the two *v*-nullclines occurs at the largest *h* value along the periodic solution. **B**. Bifurcation diagram showing how the bistability of solutions depends on *g_A_*. Arrows from panels A1 and A2 show the values of *g_A_* used to produce those simulations. The unstable fixed point (dashed blue curve) undergoes a sub-critical Hopf bifurcation as *g_A_* increases resulting in a stable fixed point and an unstable branch of periodic solutions.

Note that the bifurcation diagram is also a projection onto the *v*-*g_A_* space. In this projection, the value of *h* for the fixed point cannot be discerned. Indeed, the bifurcation diagram at *g_A_* = 0.195 μS suggests that the stable periodic solution encloses the stable fixed point (Fig. 9B). But from the projected phase plane in Figure 9A2, it appears as if the stable periodic solution encloses an unstable fixed point. In reality, neither case is true. There is a stable fixed point whose projection in the *v*-*w* phase plane occurs at the intersection of the lower *v*-nullcline and the *w*-nullcline in Figure 9A2, and which occurs for *h* = 0.45 (open circle). The intersection of the upper *v*-nullcline and the *w*-nullcline is not a fixed point, as the value of *h* is not constant along the periodic solution. Instead, it is oscillating near *h* = 0. In addition, the periodic solution has lower and upper *v* limits that are smaller and larger than the *v* value at the stable fixed point. This is why the bifurcation diagram misleadingly suggests that the stable periodic solution encloses the stable fixed point.

Figure 10 shows model predictions for how the amplitude and period of oscillations depend on either *g_NL_* (Figs. A1 and B1) or *E_NL_* (Figs. A2 and B2). The model does a reasonable job in reproducing the dependence of period on both *g_NL_* and *E_NL_* (Figs. 10A1, 10A2), at least over a range of parameters (compare to Figs. 2B and 2D, respectively).

**Figure 10.**
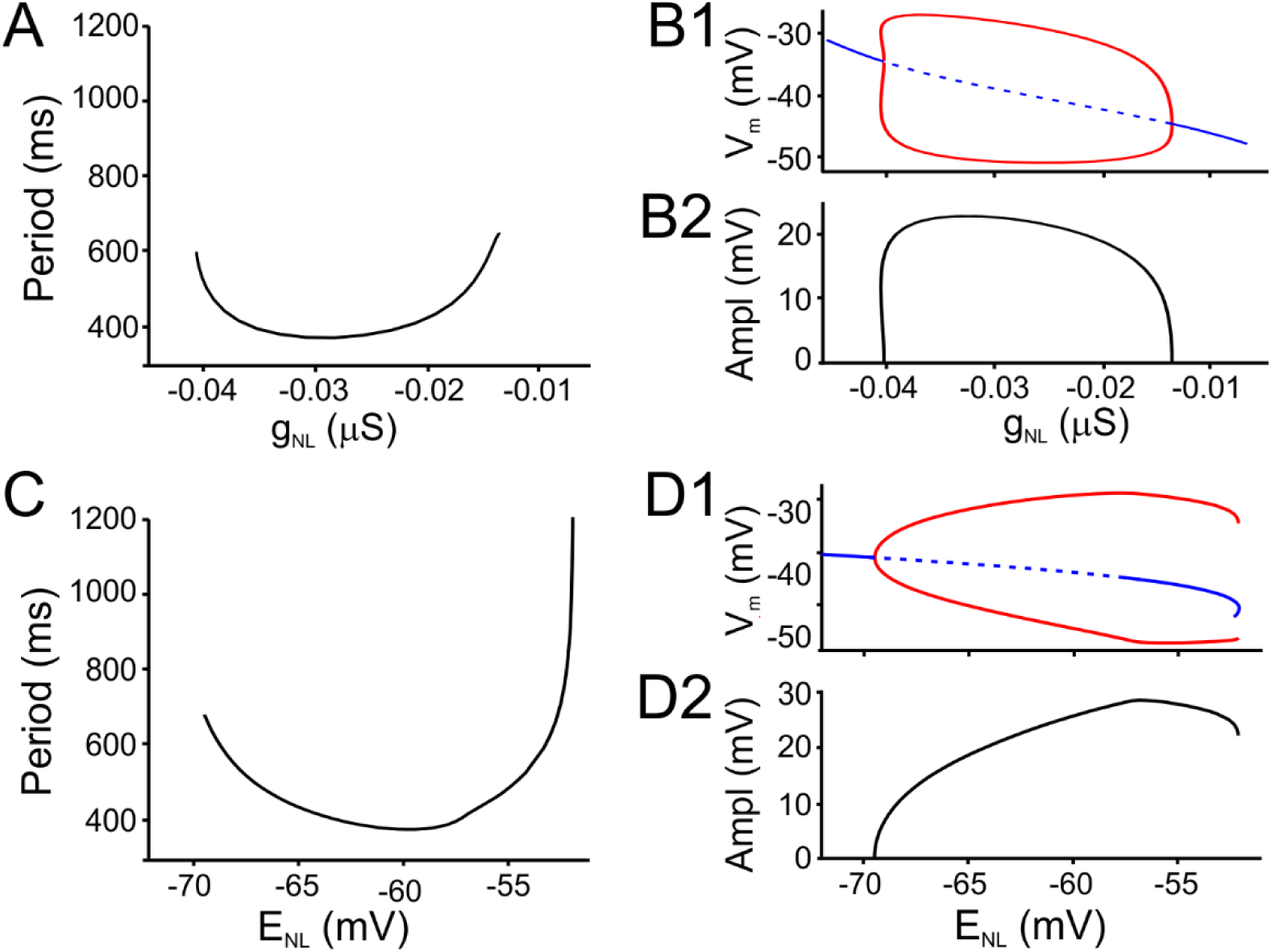
Effect of I_NL_ parameters on oscillation period and amplitude. **A, C**. Panels A and C are model analogs of Figs. 2B and 2D, showing how period depends on *g_NL_* and *E_NL_*, respectively. The simulations qualitatively match the behavior of the experimental system. **B1, D1**. Panels B1 and D1 plot the bifurcation diagrams obtained by respectively varying *g_NL_* and *E_NL_*. **B2, D2**. Panels B2 and D2 are model analogs of Figs. 2C and 2E, showing how oscillation amplitude depends on *g_NL_* and *E_NL_*, respectively. Amplitude is calculated as a difference between the minimum (bottom red) and maximum (top red) voltage for each parameter value in panels B1 and D1. The value *E_NL_* =-65 mV in Panels A and B, *g_NL_*=-0.0195 μS in Panel C and C, and *g_A_* = 0 μS in all Panels.

The dependence of the amplitude of oscillations on *g_NL_* and *E_NL_* (Figs. 10B2 and 10D2, respectively) is obtained by subtracting the upper value of voltage from the lower value (upper branch of red curve minus lower branch of red curve) in Figs. 10B1 and 10D1. The model does a poor job of describing the larger amplitude oscillations that arise in experiments for small absolute values of *g_NL_* (compare Fig. 10B2 with Fig. 2C). Mathematically this is because oscillations in the model arise due a supercritical Hopf bifurcation, whereas those that arise in the experiments may correspond to a subcritical Hopf bifurcation. The dependence of amplitude on *E_NL_* (compare Fig. 10D2 with Fig. 2E) is reasonably well described qualitatively by the model. Oscillations begin at small amplitude at low values of *E_NL_* due to a super-critical Hopf bifurcation. There is a local maximum in the amplitude at higher values of *E_NL_*, followed by a decrease in the amplitude just before oscillations cease to exist.

In sum, the simulations provide a theoretical framework to support the notion that the balance between high threshold outward currents and *I_NL_* is critical to create pre-conditions for the existence of oscillations. Once this is in place, the simulations confirm that too much *I_HTK_* (or too little total outward current) destroys this balance and thus can be used to determine which pyloric neurons can actually produce oscillations based on the *I_NL_* protocol. The model also confirms that *I_A_* does not appear to play a significant role in the generation or even modulation of oscillatory activity at least at relatively low amplitudes of the current.

## Discussion

We have shown previously (Zhao et al. 2010) that the nearly linear and negatively-sloped portion of the inward modulator-activated current (*I_MI_*) is the key element to produce oscillations in pacemaker cells of the pyloric central pattern generator. In (Bose et al. 2014), we addressed the question of what currents are minimally necessary to produce oscillatory activity in a single-cell model of a neural pattern generator. We did so by examining a simplified model of an oscillator consisting of a high threshold K^+^ current and a linear inward current that describes the behavior of this negative conductance component of *I_MI_*, and which we call the negative-conductance leak current (*I_NL_*). That work suggested, but left open, the question of how a balance between inward and distinct outward currents arises in pyloric neurons, and whether there are systematic differences in this balance between pacemaker and follower cells. Here, using both experiments and simulations, we provide evidence that pyloric neurons contain different levels of specific outward currents and that this may be the key distinction between what makes a neuron a pacemaker or a follower.

Experimentally, we find that one cell type, the PD neurons, which are considered part of the pacemaker groups of neurons in the network (Marder and Eisen 1984), are the only ones that can generate oscillations when driven by *I_NL_*: PD neurons can sustain oscillations over a large but finite range of conductances and equilibrium potentials of *I_NL_*. In contrast to the PD neurons, we find that no value of *I_NL_* conductance (or combination of *g_NL_* and *E_NL_*) can induce oscillations in any of the follower neurons. These findings are consistent with the effect the endogenous neuromodulatory peptide proctolin on the pyloric network neurons (Hooper and Marder 1987; Zhao et al. 2010). Proctolin is one of the neuromodulators that activates *I_MI_*, the current for which *I_NL_* is a simplified (linear) version. Although proctolin produces oscillations in the pyloric pacemaker neurons, it does not produce oscillations in any of the synaptically isolated pyloric follower neurons (Hooper and Marder 1987; Zhao et al. 2010).

Because pyloric neurons have the same set of ionic currents (Schulz et al. 2006; Schulz et al. 2007; Temporal et al. 2012; Temporal et al. 2014), we explored the mechanisms that preclude endogenous oscillations in the follower neurons. We found that, in follower LP and VD neurons, high-threshold outward currents (*I_HTK_*) are much too large, compared to the same currents in the PD neuron, to permit *I_NL_*-induced oscillations. However, oscillations in both of these follower neuron types could be produced if *I_HTK_* are reduced pharmacologically. These results suggests that, in order to be able to produce endogenous oscillations, a relative balance between the pacemaker current (for which *I_NL_* is a surrogate) and the counteracting outward currents needs to be maintained. Interestingly, we also observe that increasing the *I_NL_* conductance alone cannot, at least in these cells, balance the high levels of outward currents. To address why the large outward currents cannot simply be balanced by increasing the inward currents, we used a simplified mathematical model. This model confirmed that there is a finite region in the *g_NL_*-*g_HTK_* space within which oscillations are possible (Fig. 8D). The bounds on this region result from either insufficient inward current when *g_HTK_* is low (lower edge of oscillation region in Fig. 8D), or excessive leakiness of the cell when *g_HTK_* is too high, which no level of *g_NL_* can overcome to produce oscillations.

Several theoretical studies have suggested that a balance between inward and outward current is required for oscillatory activity to be generated (Doloc-Mihu and Calabrese 2014; Goldman et al. 2001; Hudson and Prinz 2010; Lamb and Calabrese 2013; Zhao and Golowasch 2012). In leech, it was shown that a close linear correlation between three currents, a leak current, a persistent K^+^ current, and a persistent Na^+^ current was required to ensure bursting oscillatory activity (Doloc-Mihu and Calabrese 2014; Lamb and Calabrese 2013). Although, in our case, the relationship between *g_K_* and *g_NL_* is not linear as in the leech studies, but rather bell-shaped and broad (Fig. 8D), those studies are consistent with ours in that relatively strict relationships must be maintained to ensure the generation of a number of features of activity, including oscillatory and bursting activity. Another example that supports the relationship between oscillatory activity and an inward/outward current balance is that of bursting pacemaker neurons in the rodent pre-Bötzinger respiratory center, in which a higher ratio of persistent Na^+^ current to leak current (I_NaP_/I_leak_) is characteristic of pacemaker neurons, while a lower ratio is typical of follower neurons (Del Negro et al. 2002).

Studies with cultured STG neurons (Haedo and Golowasch 2006; Turrigiano et al. 1995) have shown that neurons may be programmed to maintain and even restore such relationships after the loss of oscillatory activity. In these studies, recovery of rhythmic activity in cultured crab STG neurons after dissociation required the reduction of *I_HTK_*, and the increase in inward currents whose voltage dependence resembles that of *I_MI_* and *I_NL_*. These cells are capable of doing this over the course of hours to days in culture. Thus, we conclude that pacemaker activity in individual neurons involves a careful balance, not simply a linear correlation, of an inward pacemaker current with some type of current, such as *I_HTK_*, which promotes the recovery from depolarization.

In the STG, *I_HTK_* is composed of two high-threshold currents, the delayed rectifier *I_Kdr_* and the calcium-dependent *I_KCa_*. The former is known to be involved in action potential generation, while *I_KCa_* may more directly be involved in oscillatory activity (Haedo and Golowasch 2006; Soto-Trevino et al. 2005). Nevertheless, here (and earlier (Bose et al. 2014)) we showed that *I_Kdr_* is also capable of generating oscillatory activity in conjunction with *I_MI_* or *I_NL_*. In other systems too, high-threshold K^+^ currents have been shown to be essential elements of oscillatory mechanisms (e.g. Aplysia egg laying (Hermann and Erxleben 1987)) but the specific relationship with the inward currents engaged in oscillatory activity in that system have not been examined.

Interestingly, the transient *A*-current does not typically seem to be involved in pacemaker activity generation, as we confirmed in this study, even though its activation properties can be quite similar to those of high threshold K^+^ currents. In our study, *I_A_* has the same voltage-dependence of activation as *I_Kdr_*, but only *I_Kdr_* can participate in the generation of oscillatory activity (see also (Bose et al. 2014)). Thus, the differences in the kinetics and voltage dependencies of the inactivation variable are probably sufficient to determine or exclude its participation in pacemaker activity. Additionally, the transient *I_A_* was not involved in the recovery of rhythmic activity in crab cultured cells (Haedo and Golowasch 2006), consistent with the result reported here.

Our modeling results also provide the basis to interpret and understand some of our experimental results regarding the dependence of period and amplitude on parameters associated with *I_NL_*. They show that in most cases the results are qualitatively consistent over a subset of parameter values of the model. A comparison between Fig. 2B-E (experimental results) and Fig. 10 (model results) relative to how period and amplitude vary with *g_NL_* and *E_NL_*, for example, reveals a consistent relative independence of cycle period on *g_NL_* (Fig. 2B). In the model, changes in *g_NL_* simply shifted the *v*-nullcline up or down in the phase space, but left the overall balance of currents largely intact over a large range of values. As a result, neither period nor amplitude varies much (Figs. 10A and B). Variations due to changes in *E_NL_* were also consistent across both experiment and model. When *E_NL_* becomes too small, the inward effect of *I_NL_* is mitigated by the outward effect of *I_Kdr_* (their reversal potentials are too close). Alternatively when *E_NL_* is too large, the model cell is attracted to a higher level stable fixed point because the inward effect is too strong to be overcome (the driving force from *I_NL_* is too large). The model also makes predictions about how oscillations are lost through specific kinds of bifurcations as *g_NL_* or *E_NL_* are varied. It would be of interest to test if these predictions are borne out experimentally, which would provide further insight into the mechanisms of rhythm generation in pacemaker neurons similar to pyloric pacemaker neurons.

In conclusion, we observe that a coordinated balance of high-threshold outward currents and inward pacemaker currents is required for the generation of oscillatory activity. This is consistent with previous experimental and theoretical observations, but here we show both approaches confirming this in the same biological system and using a minimal model that captures the essential features of these relationships.

